# Optimal burstiness in populations of spiking neurons facilitates decoding of decreases in tonic firing

**DOI:** 10.1101/2022.04.21.488999

**Authors:** Sylvia C. L. Durian, Mark Agrios, Gregory W. Schwartz

## Abstract

A stimulus can be encoded in a population of spiking neurons through any change in the statistics of the joint spike pattern, yet we commonly summarize single-trial population activity by the summed spike rate across cells: the population peri-stimulus time histogram (pPSTH). For neurons with low baseline spike rate that encode a stimulus with a rate increase, this simplified representation works well, but for populations with high baseline rates and heterogeneous response patterns, the pPSTH has limited utility in capturing the neural representation of the stimulus. We simulated populations of spiking neurons that varied in size, baseline rate, burst statistics, and correlation, and we measured how these populations represent *decreases* (gaps) in spike rate. We introduce a different representation of the population spike pattern which we call an “information train,” and we show that it is more flexible and robust than the pPSTH in capturing stimulus information across different types of neuronal populations. In particular, we use this tool to study populations with varying levels of burstiness in their spiking statistics. We find that there is an optimal level of burstiness for gap detection that is robust to several other parameters of the population. Next, we consider this theoretical result in the context of experimental data from different types of retinal ganglion cells and determine that the baseline spike statistics of a particular, recently identified type support nearly optimal detection of both the onset and strength of a contrast step.

## Introduction

Most neurons communicate using sequences of action potentials called spike trains. Decoding information from spike trains involves extracting a signal in the face of noise, but which features of the spike train represent signal versus noise and how to represent joint spike patterns across neurons is not always clear (***Rieke et al., 1999***; ***Cessac et al., 2010***). Neurons may differ both in the way they respond to a stimulus as well as in their baseline spike patterns in the absence of a stimulus, and spike trains among a population of neurons can vary in their patterns of correlation. Many papers have examined the role of first-order statistics (mean firing rate) and correlations on the efficiency of information transmission by spike trains (***Azeredo da Silveira and Rieke, 2021***; ***Panzeri et al., 1999***; ***Ainsworth et al., 2012***; ***Ratté et al., 2013***; ***De La Rocha et al., 2007***). Higher order statistics beyond the mean and variance of the spike rate, such as temporal coding and burst coding, have also been investigated, especially in the auditory cortex (***Bowman et al., 1995; King et al., 2007***). Previous work has focused on the effect of higher-order statistics as a stimulus *response* feature but has not examined the role of higher-order baseline spike statistics in the way neural populations encode *decreases* in firing rate. Here, we develop a new analysis to represent a population response and reveal how the burstiness of spike trains affects transmission of information about firing gaps – periods of silence in spiking activity caused by a decreased firing rate.

The spike pattern of even a relatively small neural population is a distribution in high-dimensional space that is typically infeasible to sample experimentally and is likely undersampled by any down-stream decoder in the brain. Thus, finding a way to approximate this distribution and discovering which of its features are most important for a particular task, are fundamental goals in understanding neural population codes. Any change in the joint spike distribution could theoretically provide information. Sensory neurons change their firing patterns in the presence of an external stimulus, so we can measure information with respect to a known experimental variable.

Many neurons have low or zero spontaneous spike rates (***Tripathy and Gerkin, 2016***), so rate increases in the presence of a stimulus are the most common and well-studied type of information transmission (***Romo and Salinas, 2001***; ***Britten et al., 1992***; ***Gold et al., 2007***). However, the central nervous system (CNS) also contains a wide variety of tonically firing neurons, which are well suited to transmit information by *decreasing* their tonic spike rate. Examples include Purkinje cells (***Kase et al., 1980***; ***Ohtsuka and Noda, 1995***), lateral geniculate relay neurons (***McCormick and Feeser, 1990***), spinal cord neurons (***Legendre et al., 1985***), hippocampal CA1 (***Azouz et al., 1996***) and CA3 (***Raus Balind et al., 2019***) cells, and cortical pyramidal cells (***Harris et al., 2001***). Tonic firing in neurons results from a combination of the influence of synaptic networks and intrinsic electrical properties (***Llinás, 2014***; ***Destexhe and Paré, 1999***). Both factors can affect the higher-order statistics of tonic spike trains. Different neurons at the same mean rate can have spike trains which vary along the spectrum from the perfectly periodic clock-like transmission of single spikes to highly irregular bouts of bursting and silence (***Zeldenrust et al., 2018***; ***De Zeeuw et al., 2011***). The goal of the theoretical part of this study was to determine how higher-order statistics affect the encoding capacity of a neural population. Specifically, “is there an optimal level of burstiness for representing firing gaps?”

Retinal ganglion cells (RGCs), the projection neurons of the retina, offer a tractable system for both experimental and theoretical studies of information transmission in spike trains. As the sole retinal outputs, RGCs must carry all the visual information used by an animal to their downstream targets in the brain. RGC classification is more complete, particularly in mice (***Goetz et al., 2022***), than the classification of virtually any other class of neurons in the CNS. Finally, RGCs of each type form mosaics that evenly sample visual space, creating a clear relationship between the size of a visual stimulus and the number of neurons used to represent it (***Rodieck, 1998***).

Inspired by the recent discoveries of how different types of tonically firing RGCs encode contrast (***Jacoby and Schwartz, 2018***; ***Wienbar and Schwartz, 2022***), here we develop a general theoretical framework for representing information in a neural population and apply it to measure the role of burstiness in a population’s ability to encode the time and duration of a gap in firing. We propose a continuous readout of a spike train into an “information train” which tracks Shannon information (***Shannon, 1948***) over time. Critically, we show that information trains can be summed in a population in a way that captures the raw spiking activity more faithfully and completely and is more robust to population heterogeneity (in particular, heterogeneity in terms of how many cells in the population respond to the stimulus and the type of response they exhibit) than combining the underlying spike trains by standard methods such as the population peri-stimulus-time-histogram (pPSTH). We first develop a model to generate spike trains with varying levels of burstiness and correlation, then establish metrics to decode the *start time* and *duration* of a firing gap from the spike trains, and finally measure decoding performance when we vary the simulation parameters burstiness, correlation, firing rate, and population size. We assume a stimulus which depresses the firing rate in proportion to its strength; thus measuring these two characteristics of the resulting firing gap reveals both when the stimulus is presented and its strength. The goal of these simulations is to understand exactly how each of the model parameters affects a neural population’s ability to decode key pieces of information about a gap in spiking caused by a depressed firing rate, and use this to deduce how the model parameters affect ability to decode information about the stimulus.

Our principal result is a theoretical justification for why burstiness exists in some cell types: there is an optimal level of burstiness for identifying the time of stimulus presentation and stimulus strength. This result could help explain the variability in burstiness between different types of neurons. Finally, we compare our theoretical results to spike trains from recorded mouse RGCs to measure how close each type lies to optimality for each decoding task.

## Results

### A parameterized simulation of populations of tonic and bursty neurons

To study the role of higher-than-first-order statistics in the encoding of spike gaps by populations of neurons, first we needed a parameterized method to generate spike trains with systematic variation in these statistics which could accurately model experimental data. Two RGC types that our lab has studied extensively represent cases near the edges of the burstiness range. Both types have fairly similar mean tonic firing rates (between 40 and 80 Hz). However, while bursty suppressed-by-contrast (bSbC) RGCs fire in bursts of rapid activity interspersed with periods of silence, OFF sustained alpha (OFFsA) RGCs spike in more regular patterns (***Figure 1**A*) (***Wienbar and Schwartz, 2022***).

**Figure 1.**
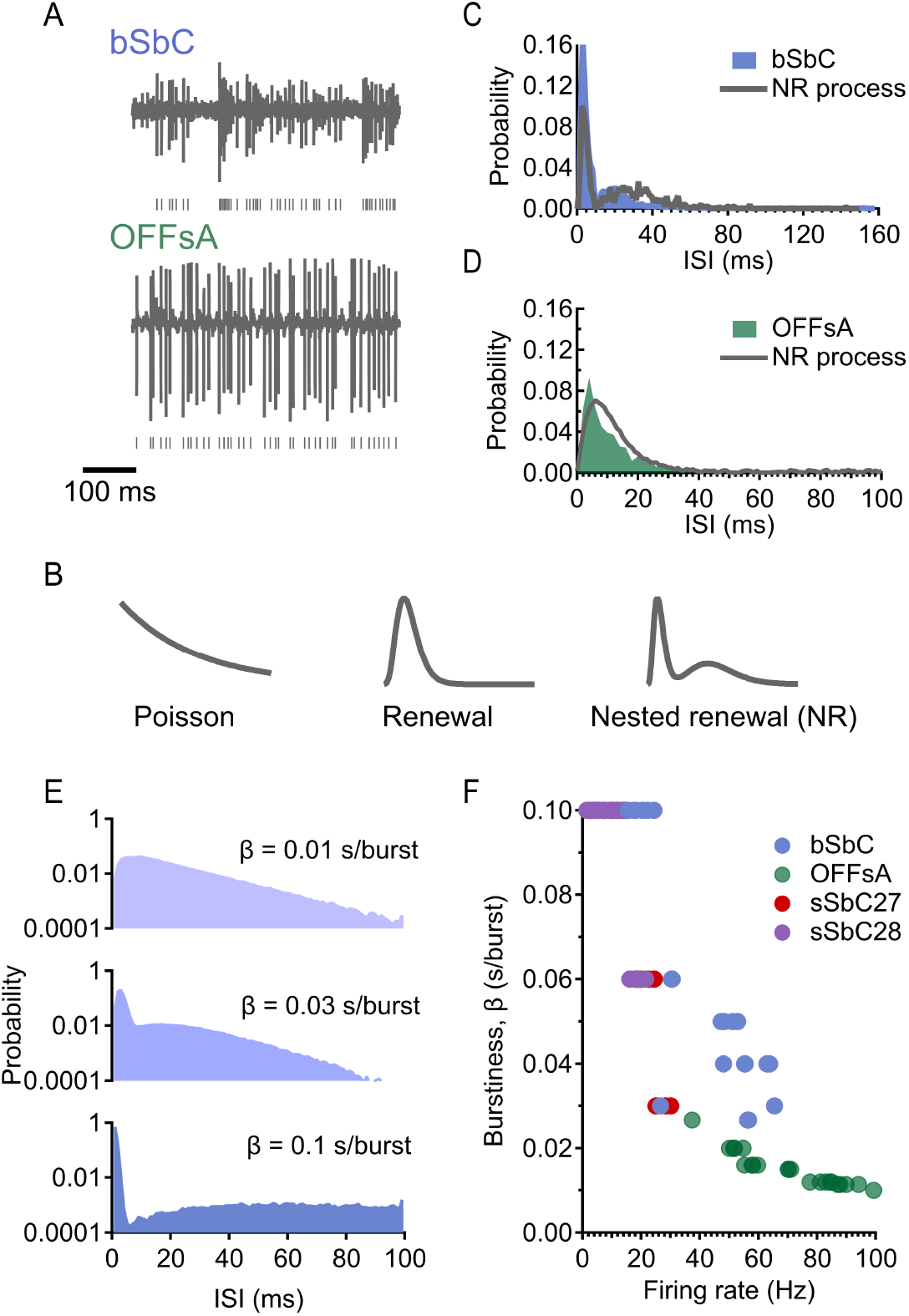
A nested renewal process can simulate bursty and tonic firing patterns with only 4 parameters. (A) Raw traces from bSbC (top) and OFFsA (bottom) in the absence of a stimulus. (B) (Left to right) ISI distributions resulting from a Poisson process (exponential), renewal process (gamma), and nested renewal process (NR; well fit by sum of gammas). (C) ISI distribution of bSbC and closest NR process match in the simulated data (as determined by a Kolmogorov-Smirov test) with parameters *κ*_1_ = 6, *λ*_1_ = 180 bursts/s, *κ*_2_ = 3, *λ*_2_ = 600 spikes/s.(D) ISI distribution of OFFsA and closest NR process match (as determined by a K-S test) with parameters *κ*_1_ = 3, *λ*_1_ = 150 bursts/s, *κ*_2_ = 3, *λ*_2_ = 500 spikes/s. (E) ISI distributions resulting from NR processes with the same firing rate of 100 Hz, ranging from regular to bursty. (Top: *κ* = 3, *λ*_1_ = 300 bursts/s, *κ*_2_ = 3, *λ*_2_ = 300 spikes/s. Middle: *κ*_1_ = 3, *λ*_1_ = 100 bursts/s, *κ*_2_ = 5, *λ*_2_ = 1500 spikes/s. Bottom: *κ*_1_ = 3, *λ*_1_ = 30 bursts/s, *κ*_2_ = 3, *λ*_2_ = 3000 spikes/s.) Bottom ISI distribution extends out to 500 ms (not shown). Note: ISI distributions are shown on a semilog scale to emphasize differences in their tails. (F) Range of firing rates and burstiness in four RGC types: bSbC and OFFsA as defined in the text, sSbC27 (sustained suppressed-by-contrast type EW27) and sSbC28 (sustained suppressed-by-contrast type EW28). Details of RGC classification can be found in ***Goetz et al**.* (***2022***) **Figure 1—figure supplement 1**. Nested renewal process ISI histograms are well fit by a sum of gammas distribution. **Figure 1—figure supplement 2**. RGC ISI histograms are well fit by a sum of gamma distribution.

A nested renewal process is a doubly stochastic point process that can simulate both bursty and tonic firing patterns with only four parameters (see **Methods**). This process builds upon one of the most commonly used methods to generate spike trains: the Poisson process. The Poisson model of spike generation assumes that the generation of spikes depends only on the instantaneous firing rate (***Vreeswijk, 2010***). Using a Poisson process to generate spikes leads to exponentially distributed interspike intervals (ISIs; ***Figure 1**B left*). One obvious limitation of Poisson spike generation, however, is its inability to model refractoriness – the period of time after a neuron spikes during which it cannot spike again. A renewal process extends the Poisson process to account for the refractory period by allowing the probability of spiking to depend on the time since the last spike as well as the instantaneous firing rate (***Vreeswijk, 2010***; ***Heeger et al., 2000***). The resulting spike train has gamma-distributed ISIs (***Figure 1**B middle*) since the refractory period does not allow for extremely short intervals between spikes.

Even though a renewal process is more physiologically accurate than a Poisson process, it still fails to model burstiness. Therefore we extended the renewal process again by nesting one renewal process inside another (***Yannaros, 1988***), similarly to how others have previously constructed doubly stochastic (although non-renewal) processes to model spikes (***Bingmer et al., 2011***). The outer renewal process parameterized by **κ**_1_ (a shape parameter) and **λ**_1_ (a rate parameter) which determines the number and placement of events that we will call “burst windows” (opportunities for a burst to occur), while the inner renewal process with parameters **κ**_2_, **λ**_2_ determines the number and placement of spikes within each burst window (see **Methods**). Both the number of burst windows and the number of spikes were randomly generated within our model, so it is important to note that a burst window could contain as few as zero spikes by chance. Therefore, a nested renewal process can flexibly simulate both regular (1 spike/burst window, as in a standard renewal process) and bursty (many spikes per burst window) firing patterns. The spike trains generated by a nested renewal process have ISIs which are well fit by a sum of gammas distribution (***Figure 1**B right* and ***Figure 1—figure Supplement 1***), where the tall, narrow left mode of the distribution corresponds to the intervals between spikes within burst windows, and the short, wide right mode of the distribution corresponds to the intervals between burst windows. Finally, a nested renewal process models activity of bSbCs and OFFsAs well; tuning its parameters results in good approximations to ISI distributions of experimentally measured bSbC (***Figure 1**C*) and OFFsA RGCs (***Figure 1**D*), since bSbCs and OFFsAs have ISI distributions which are well fit by sum of gammas as well (***Figure 1—figure Supplement 2***).

Our next goal was to quantify burstiness by a single measure. Because our method of generating spike trains necessarily places each spike within a burst window, we could not use a standard definition of burstiness, such as the total number of spikes contained within bursts divided by the total number of spikes (***Oswald et al., 2004***). Instead, we reasoned that a cell should intuitively be classified as more bursty if it contains, on average, more spikes within its burst windows. However, increasing the firing rate will automatically increase the number of spikes per burst window, so we normalized by the mean firing rate. Therefore we defined the burstiness factor, **β**, as the average number of spikes per burst window normalized by the firing rate (measured in seconds/burst window). ***Figure 1**E* illustrates the effect of different levels of burstiness on the ISI distribution for a fixed firing rate. We prefer this definition to one that thresholds the number of spikes within a certain period of time, since our definition of burstiness is easily computed from the model parameters and eliminates the need to threshold, thus reducing both the number of parameters and arbitrariness. Our quantification of burstiness is dependent upon the parameters of the nested renewal process and cannot be applied to experimental data; therefore, we matched ISI distributions from experimental data with simulated data (***Figure 1**C,D*) via a Kolmogorov-Smirov test in order to measure burstiness in spike trains from recorded RGCs (***Figure 1**F*). In addition to variable burstiness, the nested renewal process allowed us to independently modulate the firing rate and to simulate neural populations of any desired size with different levels of pairwise correlation amongst the spike trains (***Figure 2**A*; see **Methods**). For simplicity, we considered only homogeneous populations in this study: all neurons in a given population were simulated with the same model parameters (firing rate, burstiness, and pairwise correlation level). In other words, the model parameters were only varied on the population level, and not on the cell level.

**Figure 2.**
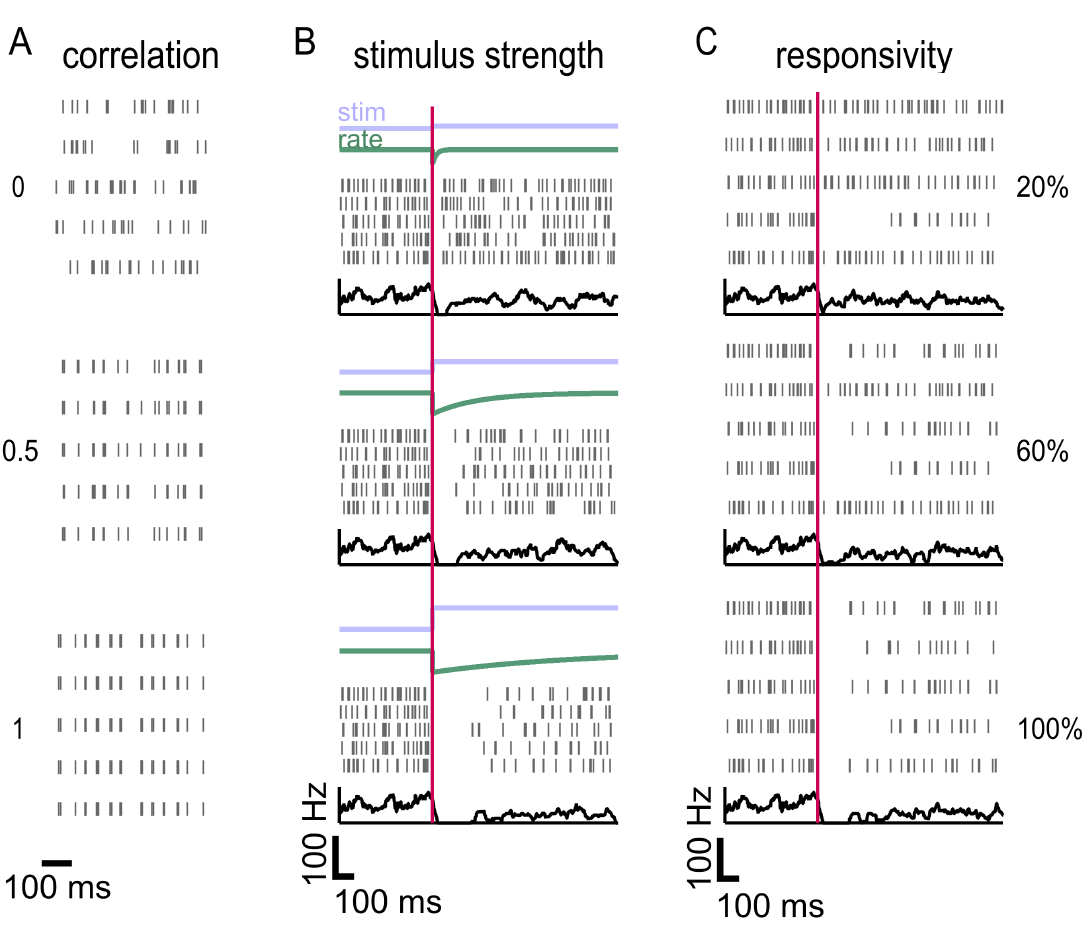
A nested renewal process can simulate populations of varying firing rate, burstiness, size, correlation and responsivity. (A) Populations of varying pairwise (neuron to neuron) correlation simulated using a nested renewal (NR) process. Rows in the rasters correspond to different neurons, not different trials. (B) Population response to varying levels of stimulus strength, modeled with a drop and subsequent rise in firing rate. The green “rate” line refers to firing rate. Population peri-stimulus-time-histogram (PSTH) is shown in black. Every neuron in each population has the same recovery time constant, but the recovery time constant varies from top to bottom (shortest for the top population and longest for the bottom) (C) Populations of varying responsivity to the stimulus. Here the recovery time constant is the same for each population and only the responsivity is varied. Population PSTH is shown in black.

We modeled a stimulus-dependent drop in firing as an instantaneous drop to a firing rate of zero followed by an exponential rise back to the baseline firing rate, controlled by a variable time constant of recovery (***Figure 2**B*). This stimulus model was chosen because it is consistent with how both bSbC and OFFsA RGCs respond to positive contrast; longer time constants of firing recovery correspond to higher contrast stimuli (***Wienbar and Schwartz, 2022***; ***Jacoby et al., 2015***). Although this stimulus model was chosen with bSbCs and OFFsAs in mind, it generalizes to any type of neuron which decreases its spontaneous firing rate proportionally to a stimulus. We also introduced heterogeneity into the population of neurons by varying *responsivity* to the stimulus, or the proportion of the population that responds to the stimulus, since this is often less than 100% in experimental studies. In other words, we made a subset of the neurons unresponsive to the stimulus by allowing them to continue spiking with baseline statistics after stimulus onset (***Figure 2**C*). Analogous to models for detection of the onset and duration of an increase in firing, this stimulus allowed us to measure performance in decoding the onset and duration of the gap in firing for each simulated population of neurons.

### The information train enables aggregation of heterogeneous spike trains

By aggregating the spike trains of the population of neurons into a one-dimensional signal over time, decoding the onset and duration of a gap in firing can be formulated as a threshold detection task: the choice of threshold determines when the decoder reports that the population is in the baseline state versus the stimulus (gap detected) state (***Figure 3**A*). The choice of threshold naturally implies a trade off between *reaction time*, the delay from stimulus onset until the threshold is crossed, and *false detection rate*, the rate at which the threshold is crossed in the absence of the stimulus. To simplify our analysis, we chose a decoder threshold for each population that achieved a fixed, low error rate of 0.1 false detections per second (we do show, however, that our results are robust to the choice of threshold in ***Figure 4—figure Supplement 1***). Thus, for the detection task, reaction time, **δ**, was the sole performance metric for the decoder. The performance metric for the duration task was more complex and is considered in the subsequent section.

**Figure 3.**
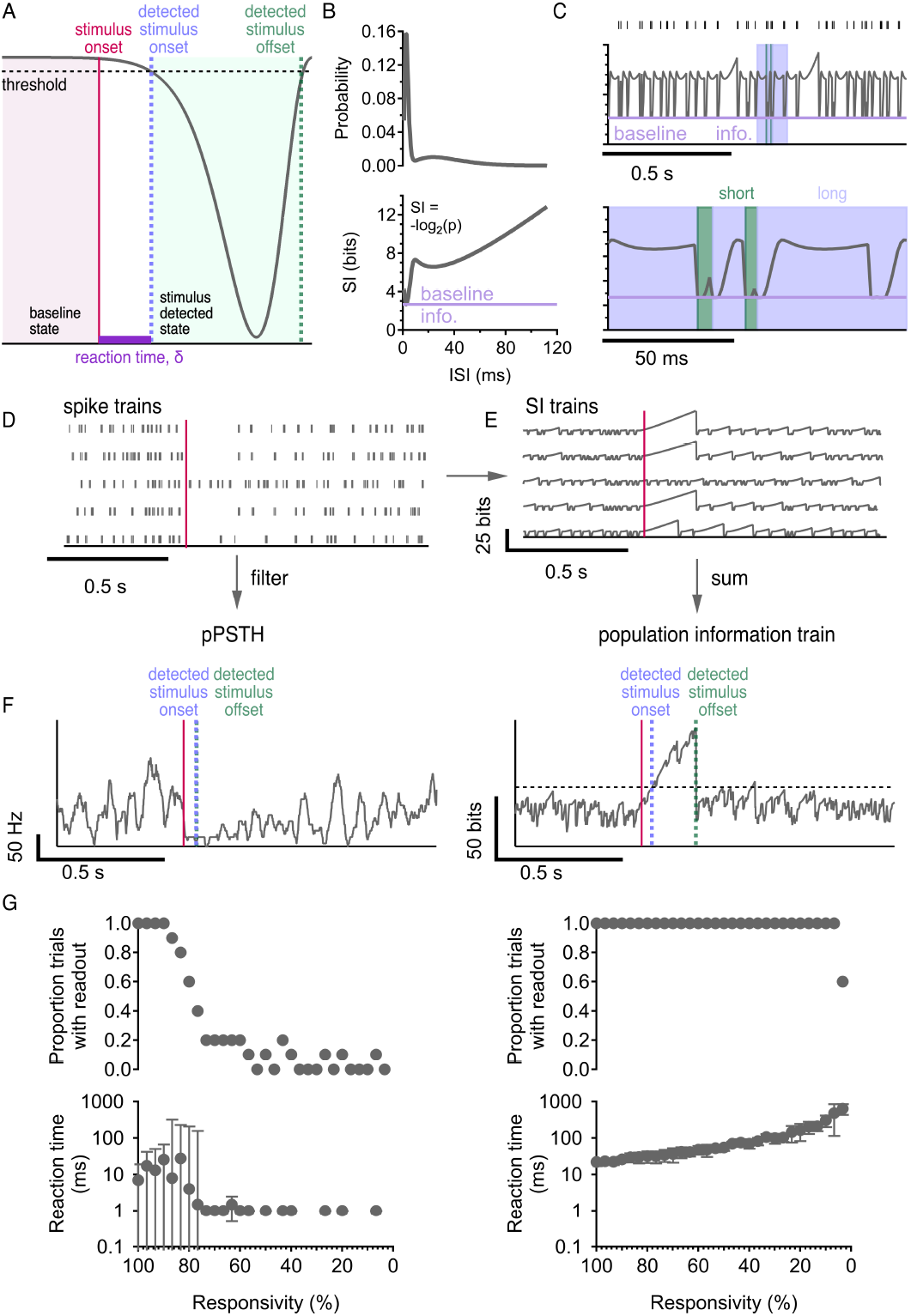
Population firing rate and population information are two plausible readout mechanisms. (A) Detecting onset and duration of a stimulus (pink) from the signal is a threshold detection task: stimulus onset (blue) is detected when the signal crosses the threshold the first time; stimulus offset (green) is detected when the signal crosses the threshold the second time. (B) ISI distribution (top) and corresponding self information (SI) curve (bottom). (C) Top: raster and corresponding single cell information train. Bottom: inset showing long ISIs (blue) and short ISIs (green). (D) Population response to a stimulus (pink) with 80% responsivity. (E) SI trains constructed from spiking activity in (D). (F) Population PSTH (left) and population information train (right) readout from activity in (D). pPSTH detected stimulus onset 50 ms after stimulus presentation; stimulus offset was detected 2 ms after stimulus onset. Information train detected stimulus onset 40 ms after stimulus presentation; stimulus offset was detected 175 ms after stimulus onset. The filter on the pPSTH and the threshold on the information train were both chosen so that there was one false detection in the prestim time (the pPSTH touched 0 once and the information train crossed the threshold once in the prestim time). Note that this is a different false detection rate than used in our study (which was 0.1 errors/s), but for demonstration purposes we wanted to show what a false detection looks like. (G) pPSTH (left) and population information train (right) readout on stimulus onset detection task as responsivity decreases in a population of 30 neurons. Top: proportion of trials with readout vs. responsivity, where readout does not exist whenever the signal fails to cross the threshold after stimulus onset. Bottom: reaction time vs. responsivity, where reaction time is the time between stimulus onset and stimulus onset detection. Error bars are interquartile range across 10 trials. **Figure 3—figure supplement 1**. Noise correlations are low.

Next, we considered the choice of the aggregation method used to combine spike trains across cells. The method most commonly used in such situations is to compute the pPSTH by simply collecting all spikes in the population and smoothing to create a continuous rate signal (***Figure 3**F right*). However, there are many properties of population activity which can carry information besides the average firing rate (***Tiesinga et al., 2008***; ***Kumbhani et al., 2007***). The pPSTH loses some of the information about the higher-order statistics of each individual spike train that could, in principle, be useful to the decoder. Therefore, we developed a new aggregation method based on the information content of the spike trains. We called the resulting signal the “information train.”

The information train method was inspired by neural self-information analysis (***Li and Tsien, 2017***). In order to build a continuous signal of the information content in a single cell over time, we started by computing a self-information (SI) curve from the ISI distribution using Shannon’s definition (***Shannon, 1948***): **SI** = −**log**_2_(**p**), where **p** is the ISI probability. The SI curve gives the information contained in every observed ISI (***Figure 3**B*). SI can be thought of as a measure of surprise; very probable ISIs correspond to small SI, while very improbable ISIs correspond to large SI. A spike train can be equally well described by its ISIs as its spike times; for each ISI observed in the spike train, we may use the SI curve to find the amount of information it contains. Neural self-information analysis then replaces each ISI in a spike train with the corresponding SI value (***Li and Tsien, 2017***), but this creates a discrete signal. Instead, our threshold detection paradigm requires a continuous signal that can be aggregated across cells.

Thus, we developed a new analysis to translate a spike train into a continuous self information train (***Figure 3**C top*). We separate ISIs into three cases. In the first case, the current ISI is of the same length as the most probable, or most commonly observed ISI for the cell. Therefore, this ISI contains the lowest possible SI, which we call the “baseline information.” The SI train always begins at baseline information, and in this case, remains there for the duration of the ISI. In the second case, the current ISI is longer than the most probable one. However, we do not know that it is longer until the point in time when it has passed the length of the most probable ISI and the cell has not yet spiked. The SI train reflects this by beginning at baseline information and staying constant for the duration of the most probable ISI, then rising according to the SI curve and stopping at the SI value indicated by the total current ISI (blue sections in ***Figure 3**C bottom*). Finally, in the third case, the current ISI is shorter than the most probable one. In this case, we know that it is shorter as soon as it ends, so the SI train stays at baseline information until the very end of the ISI, then rises instantaneously to the value indicated by the SI curve (green sections in ***Figure 3**C bottom*). At the start of each ISI, the SI train resets to baseline information, meaning that it has no memory of its history and considers subsequent ISIs independently.

For a single cell, the ISI distribution contains all the information in the spike train, so the self information train is a lossless representation. However, for multiple cells, there are joint ISI distributions to consider. Assuming independence between cells, the information contained in the population is the sum of the SI trains of each cell (see **Methods**). Independence is usually not a reasonable assumption, but experimental data from different RGC types show extremely low pair-wise noise correlations (***Figure 3—figure Supplement 1***). Correlations can also be induced by a stimulus, and stimulus correlations in populations of neurons are typically larger than noise correlations (***Schwartz et al., 2012***; ***Cafaro and Rieke, 2010***; ***Josić et al., 2009***; ***Ponce-Alvarez et al., 2013***). Our goal, however, was to study the optimal baseline spike statistics for a task in which all responsive neurons were suppressed by the stimulus simultaneously. Thus, non-independence in our population before stimulus onset and after stimulus offset (in other words, when the population was active) was only due to noise correlations. Not only are pairwise noise correlations low in different RGC types, but several studies (***Petersen et al., 2001***; ***Averbeck and Lee, 2003***; ***Oram et al., 2001***), including in the retina (***Nirenberg et al., 2001***; ***Schwartz et al., 2012***; ***Soo et al., 2011***), have shown that only a small amount of information is lost when neural responses are decoded using algorithms that ignore noise correlations – on the order of 10% of the total information. Therefore we have constructed population information trains by summing the SI trains of each cell in the population (***Figure 3**F right*) while recognizing that it is a lower bound on the total information present in the full joint distribution of the spike trains.

Although this choice of construction is certainly justifiable at the low end of correlation, where real RGC types lie, we consider correlations that go up to unity in our simulations. In the case that the population is 100% correlated, the spike trains from all the neurons will be identical, as shown in ***Figure 2**A*, and therefore the SI trains from individual neurons will all be exactly the same as well. Intuitively, the population information train should simply be equal to that of a single neuron since additional neurons offer only redundant information, but we have suggested a population information train constructed by summing all the SI trains. In fact, this is not an issue; summing identical SI trains will result in a population information train which is a scaled version of the SI train from any given individual neuron. Since we use the population information train to detect change via a threshold that is based on the false detection rate (rather than measuring the absolute value of the population information train), scaling the information train does not affect the reaction time that we measure from it. This way of constructing the population information train could break down for populations with medium pairwise correlations, but any choice of construction would have a trade off between accuracy and ease of use, and we are satisfied that this choice is easy to implement, motivated by mathematical reasoning, and most importantly works for low (which is what is actually relevant to the brain) and high correlations.

As with any decoding mechanism, the biological plausibility of the information train needs to be considered. The information train of a single neuron is based only on that neuron’s ISI distribution in the absence of a stimulus. Importantly, the information train does not need to access the whole time or the future to be built; the information can be decoded in real time if the baseline ISI distribution is known. It has been suggested that neurons can learn distributions (***Dabagia et al., 2022***), and since the baseline ISI distribution is derived from the intrinsic spiking patterns of the neuron, a readout neuron would have ample time to collect “samples” of ISIs from the neuron (or a population with identical spiking patterns within noise) and reconstruct the baseline distribution. Therefore we suggest that the information train decoding scheme is biologically plausible like a pPSTH, but captures more information.

A critical benefit of the population information train over the pPSTH is that it should automatically amplify the contribution of responsive cells in a population (which will have large SI) relative to unresponsive cells (which will have small SI) without the need to define a cell selection criterion or weighting strategy. To test this intuition, we simulated populations of 30 neurons and varied their responsivity to the stimulus. We decoded the gap onset time for all of these populations using both the pPSTH and the population information train. The pPSTH decoder was extremely sensitive to responsivity, usually failing to reach the error threshold when fewer than 80% of cells were responsive (***Figure 3**G left*). The population information train decoder approach was robust to very large fractions of unresponsive cells (***Figure 3**G right*). Subsequent analyses used the population information train decoder and 100% responsivity, but our conclusions are robust to lower responsivity (***Figure 4—figure Supplement 2***). Heterogeneity in population responsivity is just one case where the information train outperforms a pPSTH; others are considered in the Discussion. In each case, the pPSTH could potentially be modified to get around problems caused by population heterogeneity (such as a responsivity-weighted pPSTH), but the modifications would have to be different in every case. In contrast, the information train is both robust and flexible without tuning.

### There exists optimal burstiness for minimizing reaction time

Having developed a readout mechanism – the population information train – we then used it to decode the onset of a gap in firing by measuring reaction time. We were interested in the effects of multiple parameters on reaction time: burstiness, firing rate, population size, and correlation. For three of these parameters, we had an intuition for how they should affect performance based on one key insight: when trying to decode the onset of a gap, temporal precision is key to getting accurate estimates. Increasing the firing rate of a population increases temporal precision, so we expected that increasing the firing rate would improve performance (i.e. decrease reaction time). Another way to gain temporal precision is to increase the size of a population – therefore we also expected population size to have a positive effect on performance. A population which has very low cell correlations is unlikely to have many overlapping spikes from different cells, while a population with high correlation is likely to have major overlap. This implies that temporal precision, and thus performance, should decrease with correlation. In contrast to the other parameters, the intuition for how burstiness affects performance is not simple and was our central question in this part of the study.

We isolated the effect of burstiness on reaction time by holding firing rate, population size, and correlation constant and found a non-monotonic relationship (***Figure 4**A*), suggesting that for each combination of parameters there may be a different level of burstiness that minimizes reaction time. We could continue in this way to isolate the effects of each parameter by holding the other parameters constant at different values, but the number of parameters, and their ranges, makes this impractical. A more elegant approach is to try to find a unifying principle which can collapse the data. Dimensional analysis gives us the tools to identify such a unifying principle.

**Figure 4.**
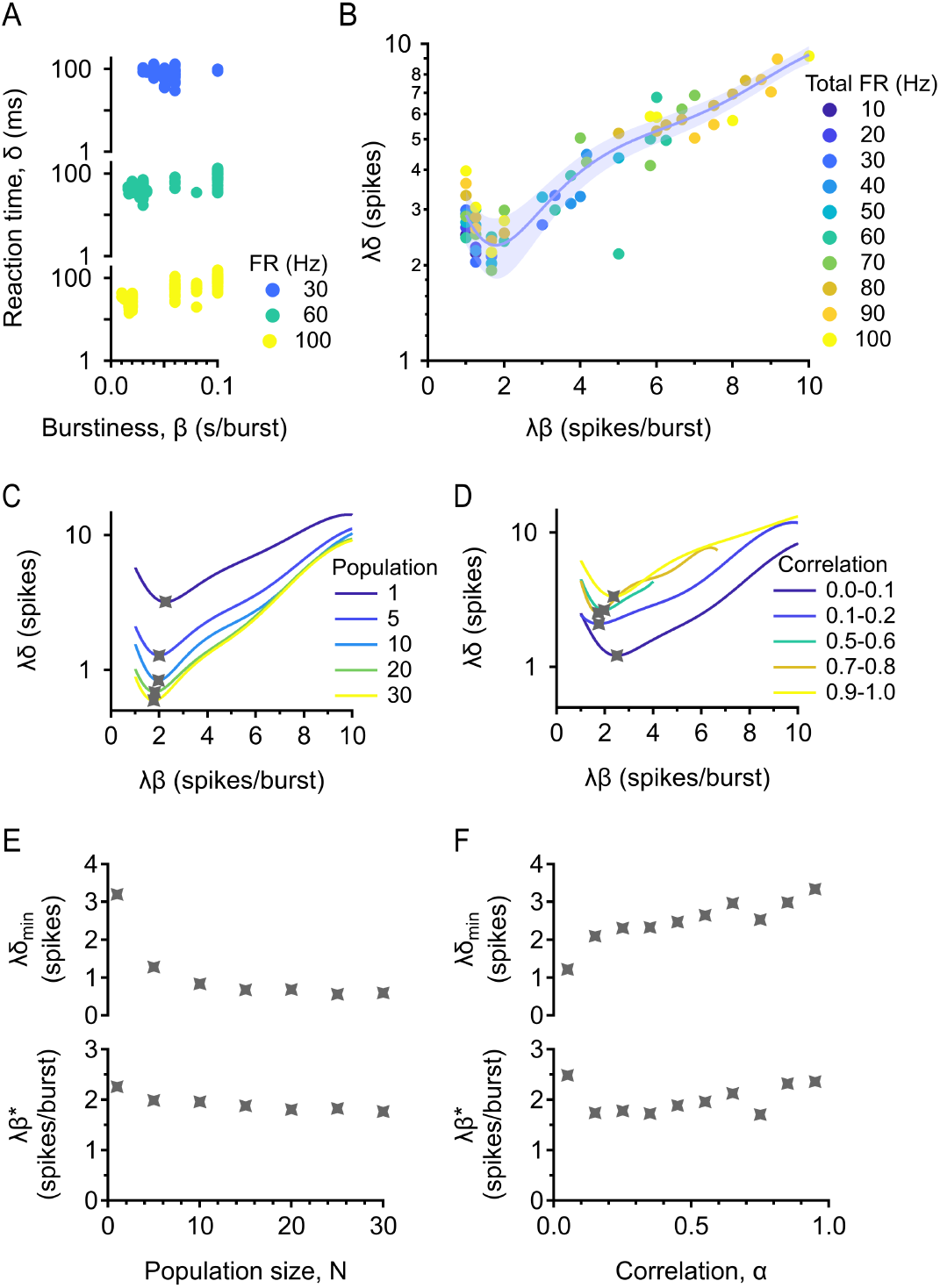
Optimal burstiness for minimizing reaction time. (A) Reaction time *δ* vs. burstiness *β* with information train readout. Population is fixed at 5 cells, correlation is fixed between 0-0.2, and total firing rate is fixed at 30, 60, and 100 Hz. (B) Dimensional analysis collapses the data for different firing rates. Dimensionless reaction time is plotted against dimensionless burstiness. Population is fixed at 5 cells and correlation is fixed between 0-0.2. Trial-averaged (median) data is shown. Error bar = standard error estimate of the data around the fit. (C) Existence of a minimum is robust across population sizes. Correlation is fixed between 0-0.2. (D) Existence of a minimum is robust across correlations. Population is fixed at 5 cells. (E) Top: minimum (dimensionless) reaction time against population size. Bottom: optimal (dimensionless) burstiness against population size. (F) Same as in (E) but correlation is varied. **Figure 4—figure supplement 1**. Results are robust to different thresholds on the information train. **Figure 4—figure supplement 2**. Results are robust to different levels of responsivity. **Figure 4—figure supplement 3**. Results are robust to different ways of constructing the population information train.

Dimensional analysis uncovers the relationships between variables by tracking their physical dimensions over calculations. Since responsivity is held constant at 100%, there are only five relevant quantities altogether: reaction time **δ**, burstiness **β**, firing rate **λ**, population size **N**, and correlation **α**. These are all either dimensionless or some transformation of the physical dimension “time,” so there is only one relevant dimension in this problem. A fundamental theorem in dimensional analysis, the Buckingham Pi Theorem (***Buckingham, 1914***), says (1) it is possible to construct exactly 5−1 = 4 independent dimensionless groups out of these five variables, and (2) those dimensionless groups are functionally related. We chose to make reaction time and burstiness dimensionless by multiplying by firing rate (brackets denote the dimension of the quantity inside and a dimensionless quanitity is said to have dimension 1), so we obtain the following four dimensionless groups:

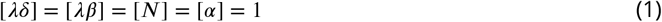

We may write any one of these dimensionless quantities as a function of the rest, but it is difficult to fit functions of three variables, so we fix population size and correlation so that the function no longer depends on them, and then we have

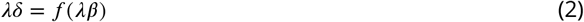

Eq (2) immediately reveals that both reaction time and burstiness are inversely proportional to the firing rate. While burstiness was defined in such a way that it must be inversely proportional to firing rate (see **Methods**), it is illuminating that reaction time is inversely proportional as well. This basic theoretical result of dimensional analysis gives us much more information than our intuition, which simply told us that reaction time should decrease with firing rate.

The practical implication of the Buckingham Pi Theorem (***Buckingham, 1914***) is that we may collapse all the data for different firing rates together, with no pattern, so we can analyze them together. The functional form of **f** is now possible to find by fitting (***Figure 4**B*). There is a clear minimum in the data, demonstrating that there is some level of burstiness which is optimal for minimizing reaction time across all firing rates. We chose a polynomial fit to describe the data, since that is a reliable way to find the minimum. We want to emphasize here that we are not claiming that the data has a polynomial form; we are only concerned with finding the minimum, and any other good fit would give the same minimum. Now we may separately vary population size and correlation (***Figure 4**C,D*), illustrating that the existence of optimal burstiness for minimizing reaction time is robust for both parameters. The x and y values of the minima of these curves completely describe how optimal burstiness and minimum reaction time depend on population size and correlation – simply dividing the x and y values of the minimum by the firing rate of the population, we obtain the exact (dimensionful) optimal burstiness and minimum reaction time. Minimum reaction time decreases with population size and increases with correlation (as predicted by our intuition), while optimal burstiness is relatively constant with both parameters, indicating that there is one level of burstiness optimal for detecting stimulus onset at any population size and correlation.

### There exists optimal burstiness for discriminating stimulus strength

Besides “when,” another fundamental question to ask about a stimulus is: how strong is it? In our model, stimulus strength corresponds to the duration of the gap in spiking because we varied stimulus strength by changing the recovery time of the firing rate (***Figure 2**B*). We measured the *gap length* in the information train (***Figure 3**E bottom right*), or the duration of time between stimulus onset and offset detection, in order to make deductions about how the length of the gap in spiking is affected by the suppressed firing rate (***Figure 5**A*). The performance metric here – the measure of how well a population can discriminate the stimulus strength – is essentially the accuracy with which the time constant of recovery is captured by the gap length measurement. Therefore the performance metric should actually be an error metric: a measure of how much error there is in the relationship between gap length and the time constant. However, it is not immediately obvious what the relationship between these two variables actually is, much less how much scatter there is around it. Dimensional analysis is again a useful tool here. There are six relevant quantities to this problem: gap length **γ**, recovery time constant **τ**, burstiness **β**, firing rate **λ**, population size **N**, and correlation **α**. Applying the Buckingham Pi Theorem (***Buckingham, 1914***) and setting all but two of the dimensionless groups (see **Methods**) to be constant (so that we obtain a function of one variable which relates the gap length and time constant of recovery), we have

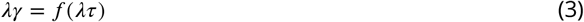

**Figure 5.**
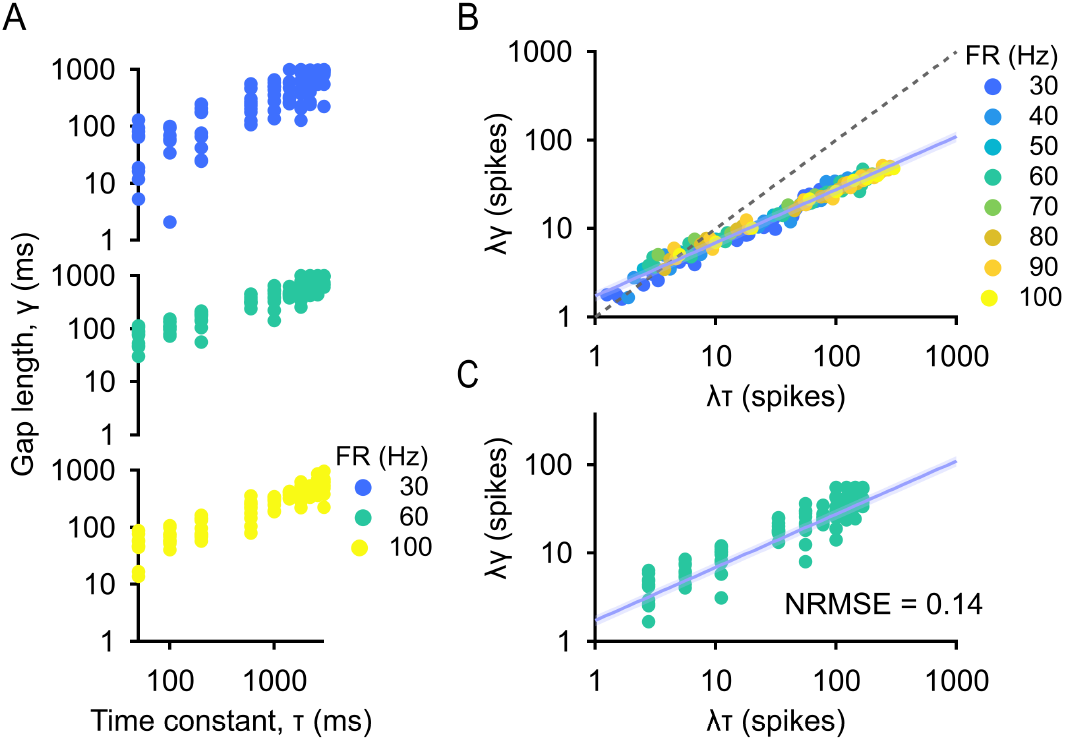
Gap length is exponentially related to the recovery time constant. (A) Gap length *γ* vs. time constant of recovery *τ*. Population is fixed at 5 cells, correlation is fixed between 0-0.2, dimensionless burstiness (*λβ*) is fixed at 3 spikes/burst, and total firing rate is fixed at 30, 60, and 100 Hz. (B) Dimensionless gap length vs dimensionless time constant. Population is 5 cells, correlation is 0-0.2, and dimensionless burstiness is 3 spikes/burst. Trial-averaged (median) data is shown. Dashed line is *y* = *x*. Error bar = standard error estimate of the data around the fit. (C) Performance metric is the scatter in the data measured with normalized root mean square error (NRMSE). Population is 5 cells, correlation is 0-0.2, and dimensionless burstiness is 3 spikes/burst, and firing rate is 60 Hz. Error bar = standard error estimate of the data around the fit.

By fitting, it is clear that there is an exponential relationship between these variables (***Figure 5**B*). Now, for each combination of parameters, for which we have several trials of gap length measurements, we chose the performance metric to be the scatter in the data around the exponential function suggested by dimensional analysis (***Figure 5**C*) which we quantified with normalized root mean square error (NRMSE; see **Methods**).

Now that we have a performance metric, we wanted to see how it depended on burstiness in particular, as well as firing rate, population size, and correlation. Once again, we could isolate the effects of each of these parameters by holding all the others constant (***Figure 6**A*), but using dimensional analysis simplifies the problem by collapsing data. The relevant quantities are the performance metric NRMSE **ϵ**, burstiness **β**, firing rate **λ**, population size **N**, and correlation **α**, so applying the Buckingham Pi Theorem (***Buckingham, 1914***) (see **Methods**) and setting population size and correlation to be constant, we have

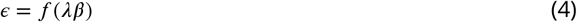

**Figure 6.**
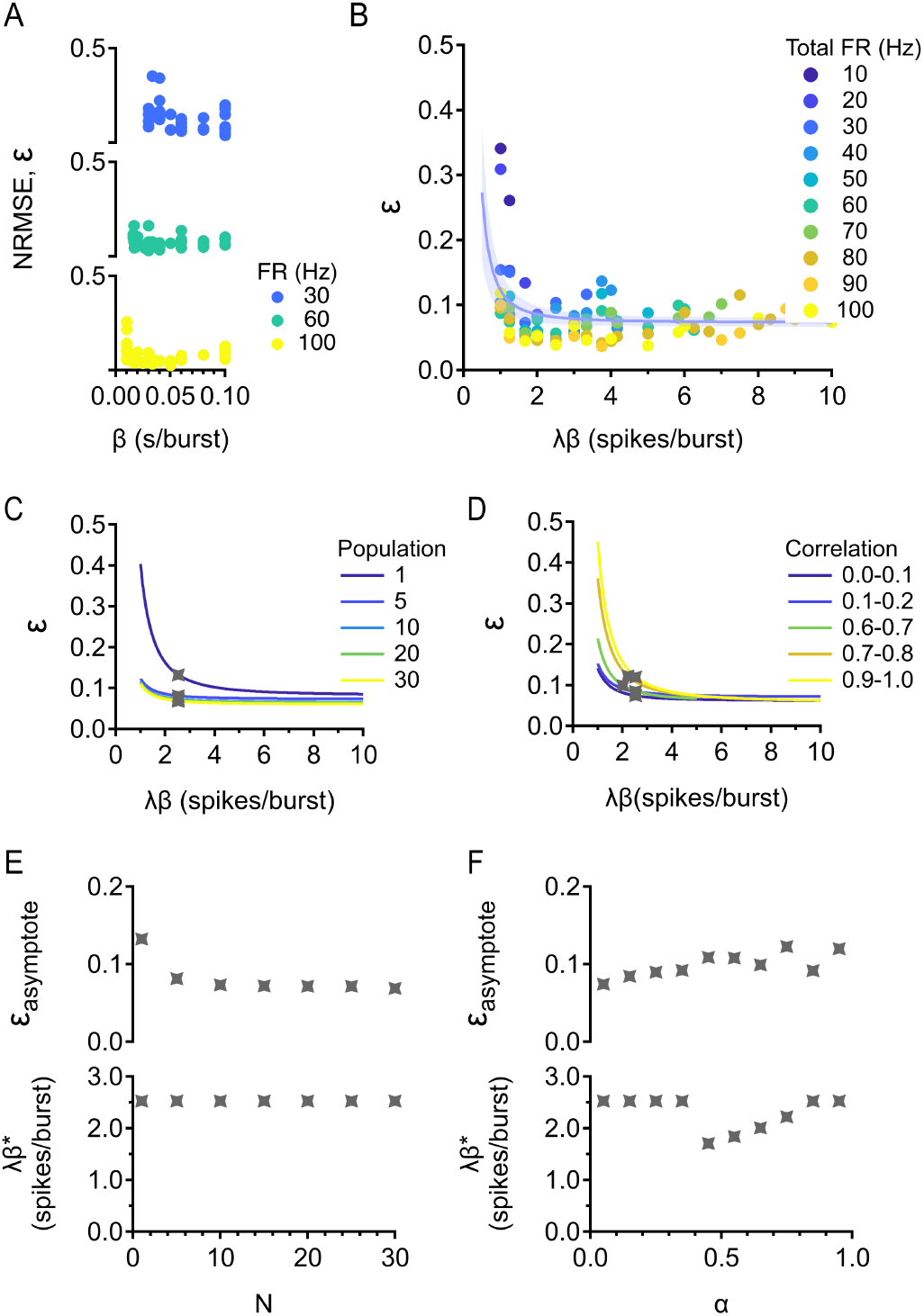
Optimal burstiness for discriminating stimulus strength. (A) NRMSE *ϵ* vs. burstiness *β*. Population is fixed at 5 cells, correlation is fixed between 0-0.2, and total firing rate is fixed at 30, 60, and 100 Hz. (B) NRMSE vs. dimensionless burstiness. Population is 5 cells and correlation is 0-0.2. Trial-averaged (median) data is shown. Error bar = standard error estimate of the data around the fit. (C) Existence of an asymptote is robust across population sizes. Correlation is fixed between 0-0.2. (D) Existence of an asymptote is robust across correlations. Population is fixed at 5 cells. (E) Top: asymptotic NRMSE against population size. Bottom: optimal (dimensionless) burstiness against population size. (F) Same as in (E) but correlation is varied.

While burstiness is still inversely proportional to the firing rate, Eq (4) reveals that NRMSE is constant with firing rate. In other words, the ability to decode the duration of a gap in firing rate does not depend on the spontaneous firing rate. Plotting reveals that NRMSE decays to a nonzero constant (***Figure 6**B*). Interestingly, there is a large range of burstiness that optimizes NRMSE, in contrast to how there was a single optimal value of burstiness for minimizing reaction time. We chose to fit an asymptotic decay function to the data (see **Methods**) in order to characterize the optimal value of NRMSE. By the nature of an asymptotic function, the fitting function continues to decay very slightly where the data has already become constant. We chose to study the “elbow” of fit because if we can describe the lowest level of burstiness that optimizes NRMSE, we will know that any burstiness larger than that will also be optimal.

Next we separately varied population size and correlation (***Figure 6**C,D*), which showed us that the existence of asymptotic NRMSE and a large range of burstiness resulting in this optimal performance was robust for both parameters. Asymptotic NRMSE decreases with population size and increases with correlation, while optimal burstiness is relatively constant with both parameters, implying that the level(s) of burstiness optimal for discriminating stimulus strength is not affected by population size or correlation.

### bSbC RGCs have close to optimal burstiness for gap detection

Having established that optimal burstiness exists for decoding the onset and duration of a gap in firing, our next question was how the baseline spike statistics of the RGC types we studied relate to this optimum. Recall that we fit functions that describe how reaction time (***Figure 4***) and NRMSE (***Figure 6***) depend on dimensionless burstiness. Dimensionless burstiness is simply bursti-ness multiplied by firing rate, so another way to represent the information in ***Figure 4***, ***Figure 6*** is as a 3 dimensional graph of reaction time (or NRMSE) vs. burstiness and firing rate (***Figure 7***). We chose to represent this information with dimensionful burstiness and firing rate instead of dimensionless burstiness because we measured those parameters directly from the data and because different cell types could potentially achieve the same dimensionless burstiness value with different combinations of firing rates and burstiness. We compared performance of different types of experimentally recorded cells by using their firing rate and burstiness to predict how quickly they would detect stimulus onset (***Figure 7**A*) and how accurately they would discriminate stimulus strength (***Figure 7**B*) according to our model. Thus the experimental data necessarily lie on the surfaces. We set the population size at 5 cells and the correlation in the physiological range of 0-0.2, but as we saw earlier, optimal burstiness is negligibly affected by both the population size and correlation so these results apply to any values of those parameters. Both bSbCs and OFFsAs are quite good at detecting a stimulus quickly, although bSbCs are closer to optimal reaction time, while both types of sustained suppressed-by-contrast RGCs are much worse at this task. However, ***Figure 7**B* suggests that bSbCs are by far the best at discriminating stimulus strength, since their burstiness puts them right on the lower bound of optimal burstiness; OFFsA RGCs do not perform nearly as well. Therefore, bSbC RGCs have spiking patterns which enable them to both react to a stimulus coming on as quickly as possible and detect the strength of that stimulus with great accuracy.

**Figure 7.**
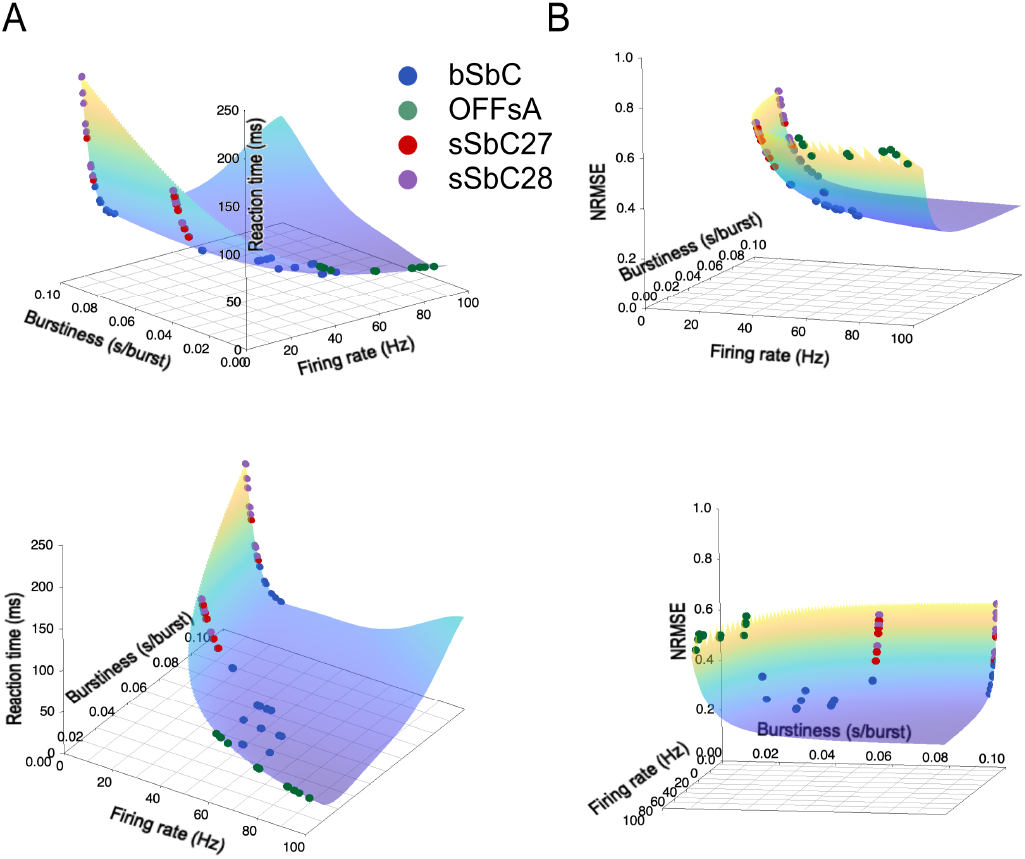
bSbCs have close to optimal burstiness. (A) 3D surface of reaction time predicted from firing rate and burstiness, as described by the function in ***Figure 4**B*. The surface is colored by elevation for clarity. The position of four RGC types on the surface is shown. Population size is 5 cells, correlation is 0-0.2. Top and bottom are different views. (B) 3D surface of NRMSE predicted from firing rate and burstiness, as described by the function in ***Figure 6**B*. The position of four RGC types on the surface is shown. Population size is 5 cells, correlation is 0-0.2. Top and bottom are different views.

## Discussion

To investigate the role of burstiness in population decoding of firing gaps, we simulated spike trains for populations of neurons using a nested renewal process. This strategy allowed us to capture the statistics of recorded spike trains from RGCs and also to vary burstiness parametrically (***Figure 1***, ***Figure 2***). We then developed a new analysis to combine spike trains across a population into an information train and demonstrated that this method is more robust to unresponsive cells than a population PSTH (***Figure 3***). Using the information trains of different simulated populations, we discovered that there is an optimal level of burstiness for detecting the onset of a firing gap that is inversely proportional to firing rate and relatively independent of population size and correlation (***Figure 4***). There is also an optimal range of burstiness for detecting gap duration that is relatively independent of all of these other parameters (***Figure 5***, ***Figure 6***). Finally, we considered the base-line spike statistics of four RGC types in the context of these theoretical results and revealed that the burst patterns of bSbC RGCs make them nearly optimal for detecting both the onset and the strength of a contrast step (***Figure 7***).

### Baseline spike statistics can be optimized for different decoding tasks

The fact that some burstiness higher than minimum is optimal for identifying the time of stimulus presentation and stimulus strength leads us to a principle that has a long history in population spike analysis (***Kepecs and Lisman, 2003***; ***Oswald et al., 2004***; ***Palmer et al., 2021***): different spiking patterns are optimal for encoding different stimulus features. We extended these results by demonstrating two concrete examples of how optimal encoding of different stimulus features depends on baseline spiking statistics. The theoretical implication of this is that different cell types may display different spiking statistics depending on the stimulus features they encode.

Earlier we provided intuition about the effects of firing rate, population size, and correlation on reaction time. The same intuition holds for performance on decoding the duration of a gap in firing, since it is also dependent on temporal precision. Now we will lay out our intuition for why there exists optimal burstiness for both tasks, which will explain why it makes sense that specific spiking patterns are optimal for encoding different stimulus features. We again attribute increased performance on these tasks to increased temporal precision. As explained earlier, a population could theoretically increase its temporal precision by increasing its firing rate or its size, or decreasing its cell correlations; however, there are physiological limits to a population’s firing rate and correlation, and the size of the population activated by a stimulus is related to the size of the stimulus (***Rodieck, 1998***). In contrast, rearranging the pattern of spikes without changing the number of spikes generated is something that a cell *could* do without expending extra energy, and if it rearranges its spiking pattern so that it has long periods of silence and short periods of rapid activity, or bursts, then it has great temporal precision within those bursts. Now imagine that that cell is part of a population with low pairwise cell correlations (***Figure 4—figure Supplement 1***) – then the bursts from different cells would be staggered in time, allowing the population as a whole to have excellent temporal precision. Therefore it is clear that increasing burstiness can improve performance. Following this line of thinking, it is also easy to see why too much burstiness would be harmful: if the population is trying to detect stimulus onset, the periods of silence between bursts could become so long that even with many cells it is very possible that the population would entirely miss the stimulus simply because no cell was in the middle of a burst when the stimulus was presented. In addition, the fact that firing rate, population size, and correlation had effects on performance that were predictable from this intuition lends further credibility to our result that there is an optimal level of burstiness for identifying the timing and strength of a stimulus.

We found that the baseline firing statistics of bSbC RGCs lie close to the optimum for performance on two gap detection tasks (***Figure 7***), but what about the other 3 high-baseline-rate RGC types we analyzed and the many lower baseline rate ones we did not analyze? It is important to consider (1) that the gap detection tasks we defined are only two out of many decoding tasks that must be performed simultaneously across the population of RGCs, and (2) that biological constraints, including the energy expenditure of maintaining a high spike rate, also likely drove the evolution of the different RGC types. Like bSbC RGCs, sSbC RGCs also signal exclusively with gaps in firing, but they do so at a substantially longer timescale (***Jacoby and Schwartz, 2018***; ***Schwartz, 2021***). Perhaps for these RGCs, the energetic benefit of maintaining a lower baseline firing rate is worth the cost of lower temporal resolution in gap detection because they represent time more coarsely than bSbC RGCs. The OFF alpha RGCs (OFFsA and OFFtA) respond to different stimuli with either increases or decreases in baseline firing (***Homann and Freed, 2017***; ***Van Wyk et al., 2009***), and OFFtA RGCs have been implicated in the detection of several specific visual features (***Münch et al., 2009***; ***Krishnamoorthy et al., 2017***; ***Wang et al., 2021***), so for these RGC types, performance in representing spike gaps needs to be balanced against metrics for their other functions.

### The information train as a way to track population-level changes in spike patterns

Our work has practical as well as theoretical implications: we proposed the information train read-out which tracks the information in a population over time, and which is more comprehensive and robust than a standard pPSTH readout. For example, if you use a pPSTH to analyze a population of direction selective cells without first finding the preferred directions of every single cell in the population, the responses from cells preferring opposite directions will cancel each other out, leading to a pPSTH which fails to reflect change due to the stimulus. The same effect will be observed if you present a light-dark edge to a population of ON or OFF cells. Of course, it is possible to remove this effect by classifying each cell’s response to the stimulus first, but that can be time consuming if the population size is large and it requires an (often arbitrary) supervised classification step. The information train will reflect a stimulus change in both of these cases even when the whole population is analyzed together. This is because any ISI length out of the ordinary (i.e. different from the most probable one) causes a positive deflection in the information train, so no matter whether a cell’s firing rate increases or decreases in response to a stimulus, the population information rises. Therefore the information train is a convenient readout mechanism to use because it does not require any assumptions to cluster cells, even when the population recording is heterogeneous. It is also easy to implement by simply fitting the observed ISI distributions in the absence of a stimulus to a gamma (many neural circuits have ISI patterns well fit by a gamma distribution (***Li et al., 2017***)) or sum of gammas distribution (depending on whether the cells in question are bursty).

### Limitations and future directions

The information train is not a complete description of the full mutual information in the population since it assumes independence. The natural extension of SI to a population is pointwise mutual information (PMI) (***Shannon, 1948***) which measures the association of single events. Much like entropy is the expected value of SI, mutual information is the expected value of PMI. In the future, a more accurate way to construct the population information train would be to use PMI: the value of the information train at time **t** is given by measuring the time since the last spike for each cell in the population and then calculating the PMI in the coincidence of those events. This is quite difficult to implement because it requires describing the joint ISI distributions. Not only is it necessary to estimate the joint ISI probabilities in the absence of a stimulus – when we can generate a lot of data but it is still difficult to estimate the joint probabilities – it is also necessary to be able to accurately extrapolate those joint ISI distributions in order to deduce the stimulus response. Namely (still assuming a stimulus which depresses firing rate), we need to be able to accurately estimate the very tails of the joint ISI distributions. We were able to do this in our study by fitting our observed ISI distributions with a sum of gammas distribution (***Figure 2—figure Supplement 2***), but it could be dangerous to first estimate the joint ISI distributions and then estimate their tail values. Even small errors in the density estimation would lead to drastically different results, since PMI (much like SI) amplifies the significance of extremely improbable events. In other words, small differences in **p** lead to large differences in **log**_2_(**p**) for low **p**. Calculating the true PMI of the population over time is an important future direction, but it has to be done carefully.

Another limitation of this study is that we have not formally seen how sensitive the information train readout is to more complex tasks/stimuli. We posit that the information train should be able to flexibly reflect any change in spiking patterns at the population level since every observed ISI different from the most common one registers a change. However, we only compared the information train to the standard pPSTH for one task; we do not know if the information train gives a more accurate or robust readout than a pPSTH for all stimuli.

Not only is there potential follow up research stemming directly from this study, such as constructing the population information train using PMI and exploring more complicated stimuli, but in addition, the information train will hopefully be used to investigate more complex statistics of spiking outside the retina, such as statistics exhibited by Purkinje cells (***Kase et al., 1980***; ***Ohtsuka and Noda, 1995***), for instance, and other variables that affect information transmission. The power of the information train is not limited to applications in neural spiking; it can be used to study any change detection task. For example, it could be applied to the problem of detecting auditory frequency change, where gaps are similarly important. The theoretical framework of the information train is general enough that it can be applied in a variety of research directions.

## Methods

### Model of spike generation

In the Poisson model of spike generation, the instantaneous firing rate is the only force which generates spikes. Assuming a constant firing rate **λ** over time, the number of spikes within a (small) interval of time Δ**t** is a Poisson distributed random variable with parameter **λ**. Using the Poisson probability density function,

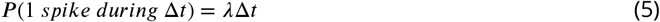

One can therefore generate a sequence of uniformly distributed random numbers {*x*_*n*_} in order to determine when to generate a spike: if *x*_*i*_ < *λ*Δ*t*, generate a spike in time bin *i*. Since the number of spikes within any interval of time is a Poisson random variable with parameter *λ*, the exponential distribution *Exp*(*λ*) describes the time between spikes, i.e., the interspike interval distribution.

A renewal process extends the Poisson process to depend on both the instantaneous firing rate and the time since the previous spike. One way to model this is to simply generate a Poisson spike train with rate parameter *λ* and delete all but every *κ^th^* spike. A gamma distribution predicts the wait time until the *κth* Poisson process event, so the ISI distribution of a renewal process generated in this way is described by *Gamma*(*κ*, *λ*). This is a natural extension of the Poisson process since if *κ* is 1, the ISI distribution is *Gamma*(1, *λ*) = *Exp*(*λ*). The average ISI length is 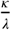 and therefore the average firing rate is 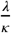.

We extended the renewal process again, nesting one renewal process inside another. The outer renewal process with parameters *κ*_1_ (a shape), *λ*_1_ (a rate) determines the number and placement of burst windows, with an average of 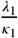 burst windows/second. We allowed to vary between 30 and 600 bursts/second, and *κ*_1_ to vary between 3 and 6 in order to obtain a range of 10-100 burst windows/second. The inner renewal process with parameters *κ*_2_, *λ*_2_ determines the number and placement of spikes within each burst window, with an average of 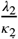 spikes/second within each burst window. We let *λ*_2_ range from 300 to 6,000 spikes/second, and *κ*_2_ range from 3 to 6. Thus the average spiking rate within a burst window ranged from 100-1000 spikes/second. We inserted spikes generated by the inside process into bursts by setting a burst length *τ*_*b*_, which we chose to be 10 ms. This results in an average of 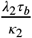 spikes/burst window, with a range of 1-10 spikes/burst window, and therefore an average firing rate

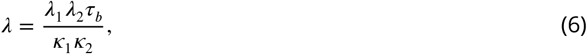

with a range of 10-100 Hz.

As a note, the burst length *τ*_*b*_ was chosen based on anecdotal observations of the typical length of burst periods in experimental recordings of bSbCs, but it is actually arbitrary. Changing the burst length would not change the ISI distributions we simulated or affect our results, because the other nested renewal process parameters can simply be changed in proportion to the burst length in order to generate the same spike train.

We varied pairwise correlations between nested renewal process spike trains by using a shared vs. random seed strategy. Just as for a Poisson process, we needed two sequences (for the inner and outer renewal processes) of uniformly distributed random numbers, {*x*_*n*_} and {*y*_*n*_}, to determine when to generate spikes (see above). We first generated two shared sequences of uniform random numbers, 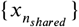 and 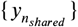, then for each cell in the population generated new independent sequences of uniformly distributed numbers, 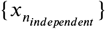 and 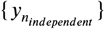 For each cell, {*x*_*n*_} was constructed by drawing from 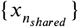 with probability *α*_*x*_ and 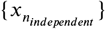 with probability 1 − *α*_*x*_. The other sequence of uniform numbers, {*y*_*n*_}, was constructed similarly with weight *α*_*y*_. The choices of *α*_*x*_ and *α*_*y*_ control the pairwise correlations of the inner and outer renewal processes, thus varying them separately varies the correlation between burst windows or the correlation between spikes within burst windows. We measured cross correlations between all pairs of spike trains in the population and averaged to obtain the overall correlation reported.

### Burstiness

#### Definition

*Burstiness β* is the average number of spikes per burst window normalized by firing rate: 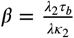.

#### Remark

Using Eq (6), the formula for burstiness can be simplified to 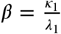. The range of burstiness is 0.01-0.1 seconds/burst window.

### Population readout mechanisms

Self information is defined as *SI* = −*log*_2_(*p*) (***Shannon, 1948***), where *p* is the ISI probability. We constructed a population information train by summing individual SI trains. This implicitly assumes that cells are independent since if independence holds, then

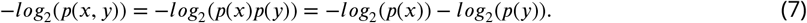

In order to measure reaction time and gap length in the information train, we set a threshold on the information train such that the error rate was 0.1 false crossings per second.

We set the threshold for the pPSTH readout mechanism at 0, and set the filter length such that the pPSTH reached 0 with a rate of 0.1 errors/second during the prestimulus time.

Population information train options B and C considered in ***Figure 4—figure Supplement 3*** were constructed as follows:

- Option B: The population information train is the SI train constructed from the ensemble spike train, or the spike train obtained by overlaying all the spike trains in the population.
- Option C: The population information train was constructed by summing the SI trains for each cell in the population, but we let ISIs shorter than the most probable one have negative deflections in each SI train, while longer ISIs still had positive deflections.

The analogous threshold (resulting in 0.1 errors/s) was placed on the information trains in options B and C.

### Dimensional analysis

We used dimensional analysis to find the relationship between gap length and the time constant of recovery (***Figure 5**A,B*). There are six relevant quantities to this problem: gap length *γ*, recovery time constant *τ*, burstiness *β*, firing rate *λ*, population size *N*, and correlation *α*. These are all either dimensionless or some transformation of “time”, so by the Buckingham Pi Theorem (***Buckingham, 1914***), we may construct 6 − 1 = 5 independent dimensionless groups related by an unknown function. We chose to make gap length, the recovery time constant, and burstiness dimensionless by multiplying by firing rate, so we have

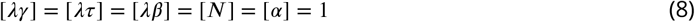

In order to obtain a function of one variable which relates gap length and the time constant, we set dimensionless burstiness, population size, and correlation to be constant and obtained Eq (3). Once we defined a performance metric (NRMSE) for the gap length decoding task (***Figure 5**C*), we used dimensional analysis again to find its dependence on burstiness and the other model parameters (***Figure 6**A,B*). The relevant quantities are the performance metric NRMSE *ϵ*, burstiness *β*, firing rate *λ*, population size *N*, and correlation *α*, so by the Buckingham Pi Theorem (***Buckingham, 1914***), we may construct 5 − 1 = 4 independent dimensionless groups related by a function. We again chose to make burstiness dimensionless by multiplying by firing rate, so we have

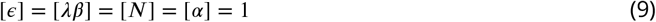

Fixing population size and correlation so that the function no longer depends on them, we obtain Eq (4).

### Fitting routines

From dimensional analysis, we obtained Eq (4). Plotting *λβ* against *ϵ* revealed that *f* is an asymptotically decaying function (***Figure 6**B*). We therefore let *f* be of the form

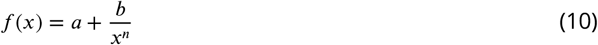

The range of *n* obtained from this fitting routine was 0.5-4 with no systematic trends, so *n* was taken as 2, which results in essentially the same good fits and reduces the number of fitting parameters to two. We also considered an exponential fit, but ultimately rejected it because the fit was not as good as the fit given by Eq (10) with fixed *n*, and it requires three fitting parameters instead of two. Goodness of fit of Eq (10) with variable *n* as well as fixed *n*, and the exponential fit, were all assessed with root mean square error (RMSE) returned by the fitting routine.

We fit the exponential relationship between the dimensionless time constant and the dimensionless gap length (***Figure 5**B,C*) by fitting a linear relationship of their logarithms. Goodness of fit was assessed with normalized root mean square error (NRMSE):

#### Definition

The NRMSE of an estimator 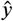 with respect to *n* observed values of a parameter *y* is the square root of the mean squared error, normalized by the mean of the measured data,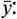 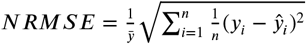

We fit ISI distributions of simulated (***Figure 1—figure Supplement 1***) and experimentally recorded data (***Figure 1—figure Supplement 2***) by minimizing the mean square error of the log of the data and log of a sum of gammas distribution, since there was a large dynamic range in the data. This effectively minimizes the percentage difference between the data and the fitting function. Goodness of fit was assessed with RMSE returned by the fitting routine.

### RGC spike recordings

RGCs were recorded in cell-attached configuration in whole-mount preparations of mouse retina as described previously (***Wienbar and Schwartz, 2022***; ***Jacoby et al., 2015***). Cell typology was determined using the methods described in (***Goetz et al., 2022***). Baseline spiking was recorded at a mean luminance of 0 and 1000 isomerizations (R*) per rod per second.

## Data and code availability

All simulated data reported in this paper will be available from a GIN database upon manuscript acceptance. Experimental data reported will be shared by the lead contact, Greg Schwartz, upon request. All code written in support of this publication will be available from a GitHub repository upon manuscript acceptance.

## Acknowledgments

We are grateful to all members of the Schwartz Lab for their feedback and support on this project. We would like to thank Stephanie Palmer and Sophia Weinbar for their comments on the manuscript, as well as Sara Solla and Ann Kennedy for their feedback on the project.

---

**Figure 1—figure supplement 1.**
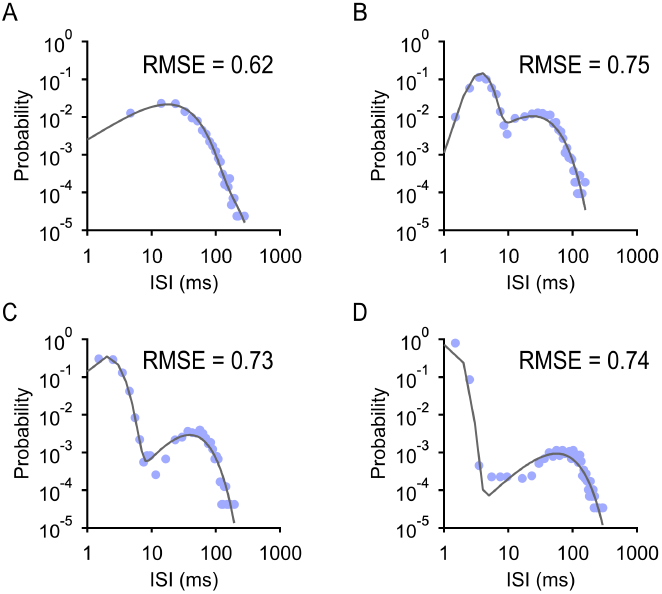
Nested renewal process ISI histograms are well fit by a sum of gammas distribution. (A) ISI distribution from NR process with parameters *κ*_1_ = 4, *λ*_1_ = 200 bursts/s, *κ*_2_ = 4, *λ*_2_ = 400 spikes/s. Firing rate *λ* = 50 Hz, burstiness *β* = 0.02 s/burst. Goodness of fit of the sum of gammas distribution is reported as root mean square error (RMSE). (B) ISI distribution from NR process with parameters *κ*_1_ = 4, *λ*_1_ = 100 bursts/s, *κ*_2_ = 5, *λ*_2_ = 1000 spikes/s. *λ* = 50 Hz, *β* = 0.04 s/burst. (C) ISI distribution from NR process with parameters *κ*_1_ = 6, *λ*_1_ = 96 bursts/s, *κ*_2_ = 3, *λ*_2_ = 1500 spikes/s. *λ* = 80 Hz, *β* = 0.06 s/burst. (D) ISI distribution from NR process with parameters *κ*_1_ = 5, *λ*_1_ = 50 bursts/s, *κ*_2_ = 6, *λ*_2_ = 6000 spikes/s. *λ* = 100 Hz, *β* = 0.1 s/burst.

**Figure 1—figure supplement 2.**
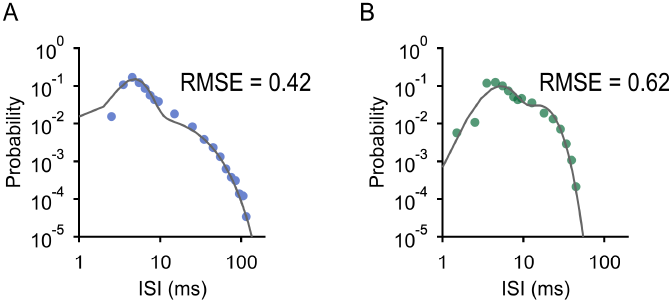
RGC ISI histograms are well fit by a sum of gamma distribution. (A) ISI distribution of experimentally recorded bSbC in **Figure 1**C with sum of gammas fit. Goodness of fit is reported as RMSE. (B) ISI distribution of experimentally recorded OFFsA in **Figure 1**D with sum of gammas fit.

**Figure 3—figure supplement 1.**
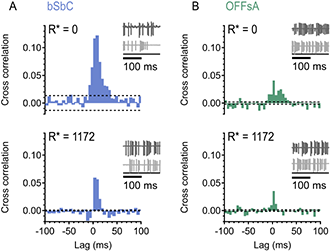
Noise correlations are low. (A) Cross correlation histograms from a pair of bSbCs. Dashed lines are null (epoch-shuffled) correlation. Paired activity was recorded at a mean luminance of 0 and 1000 isomerizations (R*) per rod per second. (B) Cross correlation histograms from a pair of OFFsAs. Dashed lines are null correlation. Paired activity was recorded at a mean luminance of 0 and 1000 isomerizations (R*) per rod per second.

**Figure 4—figure supplement 1.**
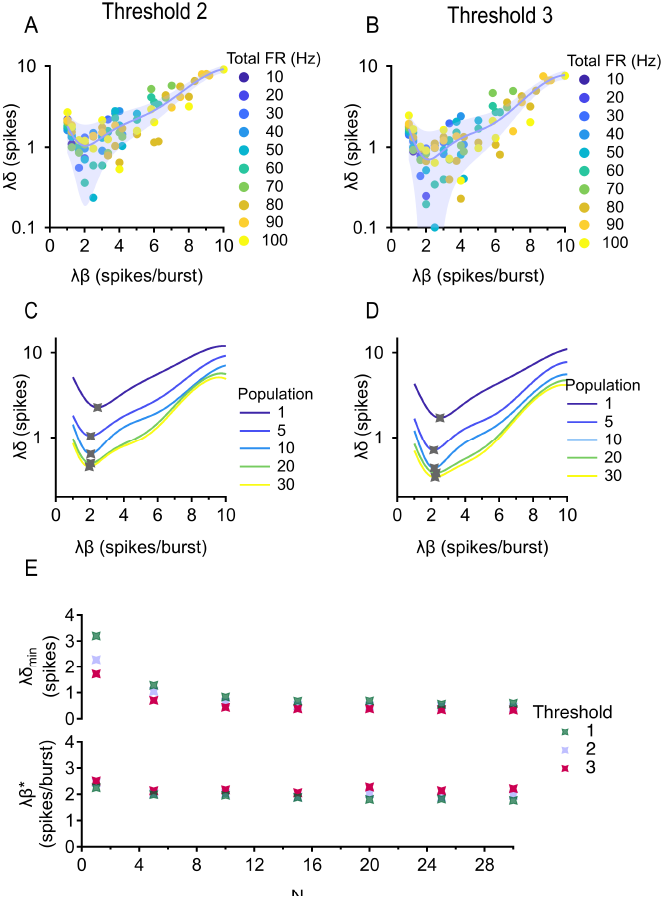
Results are robust to different thresholds on the information train. (A, B) Dimensionless reaction time measured using threshold 2 (left, 0.2 errors/s) and 3 (right, 0.4 errors/s) vs. dimensionless burstiness. Population is fixed at 5 cells and correlation is fixed between 0-0.2. Trial-averaged (median) data is shown. Error bars = standard error estimate of the data around the fits. (C, D) Existence of a minimum is robust across population sizes. Correlation is fixed between 0-0.2. (E) Top: minimum (dimensionless) reaction time against population size for thresholds 1 (0.1 errors/s), 2, and 3. Bottom: optimal (dimensionless) burstiness against population size.

**Figure 4—figure supplement 2.**
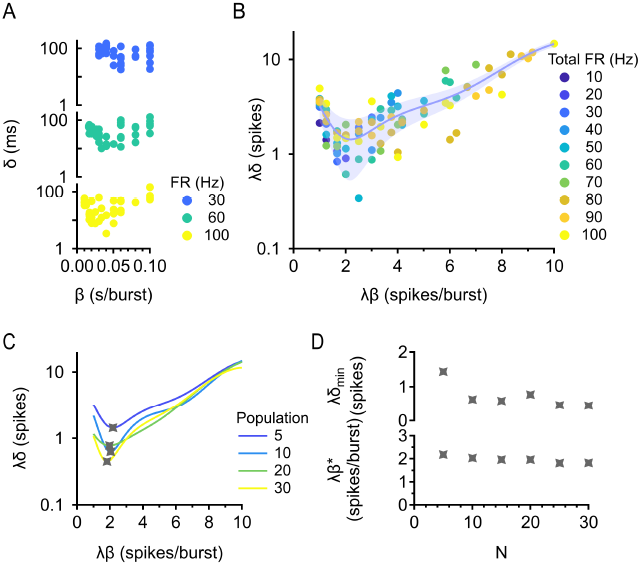
Results are robust to different levels of responsivity. (A) Reaction time *δ* vs. burstiness *β* with information train readout. Responsivity is fixed at 60%, population is fixed at 5 cells, correlation is fixed between 0-0.2, and total firing rate is fixed at 30, 60, and 100 Hz. (B) Dimensional analysis collapses the data for different firing rates. Dimensionless reaction time is plotted against dimensionless burstiness. Responsivity is fixed at 60%, population is fixed at 5 cells, and correlation is fixed between 0-0.2. Trial-averaged (median) data is shown. Error bar = standard error estimate of the data around the fit. (C) Existence of a minimum is robust across population sizes. Responsivity is fixed at 60% and correlation is fixed between 0-0.2. (D) Top: minimum (dimensionless) reaction time against population size. Bottom: optimal (dimensionless) burstiness against population size. Responsivity is fixed at 60% and correlation is fixed between 0-0.2.

**Figure 4—figure supplement 3.**
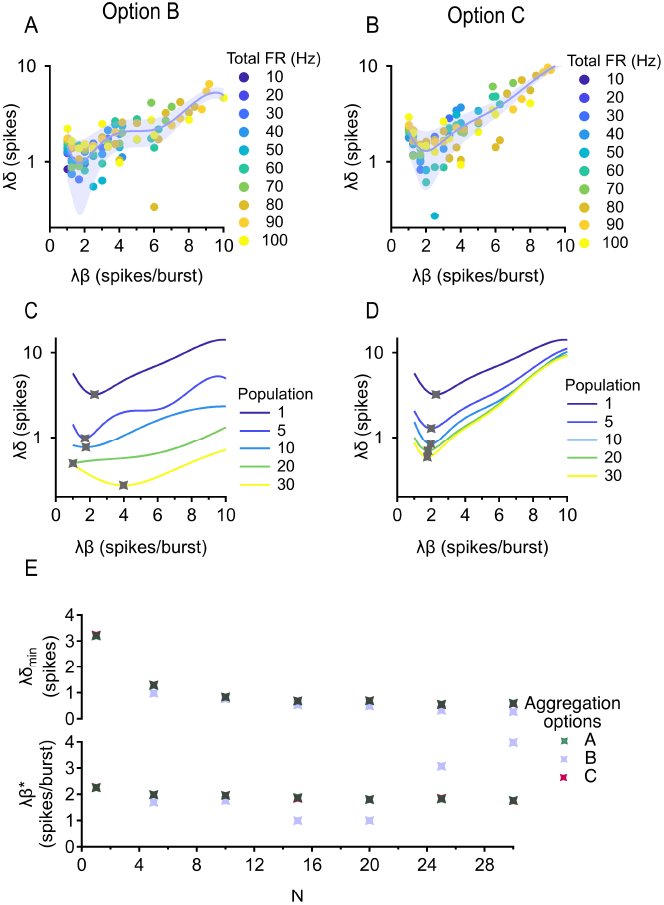
Results are robust to different ways of constructing the population information train. (A, B) Dimensionless reaction time measured using aggregation options B (left) and C (right) vs. dimensionless burstiness. Population is fixed at 5 cells and correlation is fixed between 0-0.2. Trial-averaged (median) data is shown. Error bars = standard error estimate of the data around the fits. (C, D) Existence of a minimum is robust across population sizes. Correlation is fixed between 0-0.2. (E) Top: minimum (dimensionless) reaction time against population size for aggregation options A, B, and C. Bottom: optimal (dimensionless) burstiness against population size.

## References

Ainsworth M, Lee S, Cunningham MO, Traub RD, Kopell NJ, Whittington MA. Rates and rhythms: a synergistic view of frequency and temporal coding in neuronal networks. Neuron. 2012; 75(4):572–583.

Averbeck BB, Lee D. Neural noise and movement-related codes in the macaque supplementary motor area. Journal of Neuroscience. 2003; 23(20):7630–7641.

Azouz R, Jensen MS, Yaari Y. Ionic basis of spike after-depolarization and burst generation in adult rat hippocampal CA1 pyramidal cells. The Journal of physiology. 1996; 492(1):211–223.

Bingmer M, Schiemann J, Roeper J, Schneider G. Measuring burstiness and regularity in oscillatory spike trains. Journal of neuroscience methods. 2011; 201(2):426–437.

Bowman DM, Eggermont J, Smith GM. Effect of stimulation on burst firing in cat primary auditory cortex. Journal of neurophysiology. 1995; 74(5):1841–1855.

Britten KH, Shadlen MN, Newsome WT, Movshon JA. The analysis of visual motion: a comparison of neuronal and psychophysical performance. Journal of Neuroscience. 1992; 12(12):4745–4765.

Buckingham E. On physically similar systems; illustrations of the use of dimensional equations. Physical review. 1914; 4(4):345.

Cafaro J, Rieke F. Noise correlations improve response fidelity and stimulus encoding. Nature. 2010; 468(7326):964–967.

Cessac B, Paugam-Moisy H, Viéville T. Overview of facts and issues about neural coding by spikes. Journal of Physiology-Paris. 2010; 104(1-2):5–18.

Dabagia M, Vempala SS, Papadimitriou C. Assemblies of neurons learn to classify well-separated distributions. In: Conference on Learning Theory PMLR; 2022. p. 3685–3717.

De La Rocha J, Doiron B, Shea-Brown E, Josić K, Reyes A. Correlation between neural spike trains increases with firing rate. Nature. 2007; 448(7155):802–806.

De Zeeuw CI, Hoebeek FE, Bosman LW, Schonewille M, Witter L, Koekkoek SK. Spatiotemporal firing patterns in the cerebellum. Nature Reviews Neuroscience. 2011; 12(6):327–344.

Destexhe A, Paré D. Impact of network activity on the integrative properties of neocortical pyramidal neurons in vivo. Journal of neurophysiology. 1999; 81(4):1531–1547.

Goetz J, Jessen ZF, Jacobi A, Mani A, Cooler S, Greer D, Kadri S, Segal J, Shekhar K, Sanes JR, et al. Unified classification of mouse retinal ganglion cells using function, morphology, and gene expression. Cell reports. 2022; 40(2):111040.

Gold JI, Shadlen MN, et al. The neural basis of decision making. Annual review of neuroscience. 2007; 30(1):535–574.

Harris KD, Hirase H, Leinekugel X, Henze DA, Buzsáki G. Temporal interaction between single spikes and complex spike bursts in hippocampal pyramidal cells. Neuron. 2001; 32(1):141–149.

Heeger D, et al. Poisson model of spike generation. Handout, University of Standford. 2000; 5(1-13):76.

Homann J, Freed MA. A mammalian retinal ganglion cell implements a neuronal computation that maximizes the SNR of its postsynaptic currents. Journal of Neuroscience. 2017; 37(6):1468–1478.

Jacoby J, Schwartz GW. Typology and circuitry of suppressed-by-contrast retinal ganglion cells. Frontiers in Cellular Neuroscience. 2018; 12:269.

Jacoby J, Zhu Y, DeVries SH, Schwartz GW. An amacrine cell circuit for signaling steady illumination in the retina. Cell reports. 2015; 13(12):2663–2670.

Josić K, Shea-Brown E, Doiron B, de la Rocha J. Stimulus-dependent correlations and population codes. Neural computation. 2009; 21(10):2774–2804.

Kase M, Miller DC, Noda H. Discharges of Purkinje cells and mossy fibres in the cerebellar vermis of the monkey during saccadic eye movements and fixation. The Journal of physiology. 1980; 300(1):539–555.

Kepecs A, Lisman J. Information encoding and computation with spikes and bursts. Network: Computation in neural systems. 2003; 14(1):103.

King AJ, Bajo VM, Bizley JK, Campbell RA, Nodal FR, Schulz AL, Schnupp JW. Physiological and behavioral studies of spatial coding in the auditory cortex. Hearing research. 2007; 229(1-2):106–115.

Krishnamoorthy V, Weick M, Gollisch T. Sensitivity to image recurrence across eye-movement-like image transitions through local serial inhibition in the retina. Elife. 2017; 6:e22431.

Kumbhani RD, Nolt MJ, Palmer LA. Precision, reliability, and information-theoretic analysis of visual thalamo-cortical neurons. Journal of neurophysiology. 2007; 98(5):2647–2663.

Legendre P, McKenzie J, Dupouy B, Vincent J. Evidence for bursting pacemaker neurones in cultured spinal cord cells. Neuroscience. 1985; 16(4):753–767.

Li M, Xie K, Kuang H, Liu J, Wang D, Fox G, et al. Spike-timing patterns conform to gamma distribution with regional and cell type-specific characteristics. Biorxiv. 2017; 145813(10.1101):145813.

Li M, Tsien JZ. Neural code—neural self-information theory on how cell-assembly code rises from spike time and neuronal variability. Frontiers in cellular neuroscience. 2017; 11:236.

Llinás RR. Intrinsic electrical properties of mammalian neurons and CNS function: a historical perspective. Frontiers in cellular neuroscience. 2014; 8:320.

McCormick D, Feeser H. Functional implications of burst firing and single spike activity in lateral geniculate relay neurons. Neuroscience. 1990; 39(1):103–113.

Münch TA, Da Silveira RA, Siegert S, Viney TJ, Awatramani GB, Roska B. Approach sensitivity in the retina processed by a multifunctional neural circuit. Nature neuroscience. 2009; 12(10):1308–1316.

Nirenberg S, Carcieri SM, Jacobs AL, Latham PE. Retinal ganglion cells act largely as independent encoders. Nature. 2001; 411(6838):698–701.

Ohtsuka K, Noda H. Discharge properties of Purkinje cells in the oculomotor vermis during visually guided saccades in the macaque monkey. Journal of Neurophysiology. 1995; 74(5):1828–1840.

Oram MW, Hatsopoulos NG, Richmond BJ, Donoghue JP. Excess synchrony in motor cortical neurons provides redundant direction information with that from coarse temporal measures. Journal of neurophysiology. 2001; 86(4):1700–1716.

Oswald AMM, Chacron MJ, Doiron B, Bastian J, Maler L. Parallel processing of sensory input by bursts and isolated spikes. Journal of Neuroscience. 2004; 24(18):4351–4362.

Palmer SE, Wright B, Doupe AJ, Kao MH. Variable but not random: temporal pattern coding in a songbird brain area necessary for song modification. Journal of Neurophysiology. 2021; 125(2):540–555.

Panzeri S, Schultz SR, Treves A, Rolls ET. Correlations and the encoding of information in the nervous system. Proceedings of the Royal Society of London Series B: Biological Sciences. 1999; 266(1423):1001–1012.

Petersen RS, Panzeri S, Diamond ME. Population coding of stimulus location in rat somatosensory cortex. Neuron. 2001; 32(3):503–514.

Ponce-Alvarez A, Thiele A, Albright TD, Stoner GR, Deco G. Stimulus-dependent variability and noise correlations in cortical MT neurons. Proceedings of the National Academy of Sciences. 2013; 110(32):13162–13167.

Ratté S, Hong S, De Schutter E, Prescott SA. Impact of neuronal properties on network coding: roles of spike initiation dynamics and robust synchrony transfer. Neuron. 2013; 78(5):758–772.

Raus Balind S, Magó Á, Ahmadi M, Kis N, Varga-Németh Z, Lőrincz A, Makara JK. Diverse synaptic and dendritic mechanisms of complex spike burst generation in hippocampal CA3 pyramidal cells. Nature communications. 2019; 10(1):1–15.

Rieke F, Warland D, Van Steveninck RdR, Bialek W. Spikes: exploring the neural code. MIT press; 1999.

Rodieck RW. The first steps in seeing, vol. 1. Sinauer Associates Sunderland, MA; 1998.

Romo R, Salinas E. Touch and go: decision-making mechanisms in somatosensation. Annual review of neuro-science. 2001; 24(1):107–137.

Schwartz G. Retinal Computation. Academic Press; 2021.

Schwartz G, Macke J, Amodei D, Tang H, Berry MJ. Low error discrimination using a correlated population code. Journal of neurophysiology. 2012; 108(4):1069–1088.

Shannon CE. A mathematical theory of communication. The Bell system technical journal. 1948; 27(3):379–423.

Azeredo da Silveira R, Rieke F. The geometry of information coding in correlated neural populations. Annual Review of Neuroscience. 2021; 44:403–424.

Soo FS, Schwartz GW, Sadeghi K, Berry MJ. Fine spatial information represented in a population of retinal ganglion cells. Journal of Neuroscience. 2011; 31(6):2145–2155.

Tiesinga P, Fellous JM, Sejnowski TJ. Regulation of spike timing in visual cortical circuits. Nature reviews neuro-science. 2008; 9(2):97–107.

Tripathy S, Gerkin R, Spontaneous firing rate; 2016. https://neuroelectro.org/ephys_prop/18/.

Van Wyk M, Wässle H, Taylor WR. Receptive field properties of ON-and OFF-ganglion cells in the mouse retina. Visual neuroscience. 2009; 26(3):297–308.

Vreeswijk Cv. Stochastic models of spike trains. Analysis of parallel spike trains. 2010; p. 3–20.

Wang F, Li E, De L, Wu Q, Zhang Y. OFF-transient alpha RGCs mediate looming triggered innate defensive response. Current Biology. 2021; 31(11):2263–2273.

Wienbar S, Schwartz GW. Differences in spike generation instead of synaptic inputs determine the feature selectivity of two retinal cell types. Neuron. 2022;.

Yannaros N. On Cox processes and gamma renewal processes. Journal of applied probability. 1988; 25(2):423–427.

Zeldenrust F, Wadman WJ, Englitz B. Neural coding with bursts—current state and future perspectives. Frontiers in computational neuroscience. 2018; 12:48.

